# Schistosome exposure and diet induced effects on candidate immune gene expression in an African snail vector

**DOI:** 10.1101/2025.05.06.651976

**Authors:** Tom Pennance, Johannie Spaan, Yijia Xiong, Alexa Churan, Angela Loczi-Storm, Daisy Ward, Tasneem Islam, Ashleigh Calcote, Erica Fuller, Brent Marsonette, Maurice Odiere, Michelle Steinauer

## Abstract

*Schistosoma mansoni* is a parasitic helminth that is vectored through freshwater snails. While the anti-schistosome defense of the South American snail, *Biomphalaria glabrata*, is well studied, little is known about the immune response of the African snail, *Biomphalaria sudanica*. We measured expression of five candidate immune genes in *B. sudanica* 8, 24, and 72 hours post-exposure to *S. mansoni* using reverse transcription quantitative PCR. Expression patterns of resistant snails were compared to susceptible snails and those sham exposed. We also assessed how diet (lettuce vs. pellet) affected expression of three genes, given prior findings that pellet-fed snails were more susceptible to *S. mansoni*. Results indicated that resistant snails constitutively expressed higher levels of superoxide dismutase 1 (SOD1) than susceptible snails, consistent with expression patterns of resistant *B. glabrata*. Parasite-induced expression occurred at 8 hours in SOD1, biomphalysin, thioester protein 1 (TEP1), and granulin (GRN); however, for biomphalysin and TEP1, induced expression was only detected for susceptible snails. At 24 hours, biomphalysin expression increased in exposed resistant snails, and at 72 hours, all exposed snails decreased biomphalysin expression compared to controls. Parasite-induced expression of SOD1, biomphalysin, TEP1, and GRN supports the hypothesis that these genes play a role in *B. sudanica* anti-schistosome defense, however increased expression does not necessarily yield clearance of *S. mansoni*. SOD1 expression was higher in lettuce-fed snails at 8 and 24 hours, consistent with their greater resistance. Together, these results demonstrate the conserved and unique aspects of the *B. sudanica* anti-schistosome response.

## Introduction

Schistosomiasis is a neglected tropical disease caused by infectious blood flukes that are transmitted via freshwater intermediate snail hosts. Current public health measures aimed at reducing the burden of schistosomiasis in endemic regions consist primarily of mass drug administration of praziquantel to vulnerable populations (WHO 2022; Kokaliaris et al. 2022). Approaches aimed at reducing transmission of the disease, such as molluscicides or habitat modification to reduce or eliminate snail vector populations, are ineffective in large bodies of water, lack specificity to snails, and are costly (King et al. 2015; Garba Djirmay et al. 2024). Consequentally, snail control is seldom utilized in contemporary control programs, despite its historical importance in preventing schistosomiasis in certain endemic regions (King and Bertsch 2015). A thorough understanding of the intermediate host’s means of resistance may contribute to more effective and ecologically sustainable practices of curtailing transmission.

While snails are often referred to as either resistant or susceptible to schistosomes, the relationship between snails and schistosomes is more accurately characterized as a compatibility polymorphism in which successful infections are dependent on both the host and parasite involved (Basch 1975; Richards and Shade 1987; Webster and Davies 2001; Théron and Coustau 2005; Mitta et al. 2017; Spaan et al. 2023). Most knowledge about the snail immune response to schistosomes comes from studies comparing responses of genetic lines of the South American species *Biomphalaria glabrata* that are either compatible or incompatible with a specific line of *Schistosoma mansoni*. These studies have revealed that the innate immune response of *B. glabrata* to *S. mansoni* infection involves both humoral and cell-mediated components.

In the case of an incompatible snail-parasite combination, humoral factors such as fibrinogen related proteins (FREPs) (Adema et al. 1997; Mitta et al. 2017; Pila et al. 2017) and thioester proteins (TEP) (Portet et al. 2018; Marquez et al. 2022) bind to the invading parasite and recruit biomphalysin, an aerolysin pore forming toxin, to attack the parasite (Galinier et al. 2013; Li et al. 2020; Pinaud et al. 2021).

Schistosome invasion also stimulates a cellular immune response. During this process, hemocytes, the primary immune cells, differentiate into distinct subtypes that appear to have varying roles in the immune response (Li et al. 2022; Pichon et al. 2022). An endogenous growth factor granulin (GRN) drives differentiation and proliferation of granulocytic hemocytes (Pila et al. 2016a; Hambrook et al. 2020; Bowhay and Hanington 2024). These hemocytes are recruited to the invading sporocyst, where they encapsulate the parasite and release cytotoxic reactive oxygen and nitrogen species to destroy it (Hahn et al. 2001a, b). The successful clearance of schistosomes by snails is thought to be initiated by parasite recognition (Pila et al. 2017). Several potential pathogen recognition receptors (PRRs) have been identified in *B. glabrata*, including a toll-like receptor (TLR) that is upregulated in resistant snails following exposure to schistosomes (Pila et al. 2016b).

While many of the aspects of the *B. glabrata* immune response are likely conserved among species of *Biomphalaria*, differences can be expected due to the unique evolutionary histories and co-evolutionary histories of these snails with pathogens. This is particularly relevant for species endemic to sub-Saharan Africa, which 1) diverged from a common ancestor with *B. glabrata* between ∼1.8 to 5 MYA (Woodruff and Mulvey 1997; Campbell et al. 2000; DeJong et al. 2001; Bu et al. 2023) and 2) share the longest evolutionary history with *S. mansoni*, dating back to the parasites origin in east Africa around ∼0.4 MYA (Morgan et al. 2005; Crellen et al. 2016; Platt et al. 2022).

Our study aimed to describe the expression patterns of the orthologs of five putative anti-schistosome defense genes in *B. sudanica s*.*l*., following exposure to *S. mansoni. Biomphalaria sudanica* is the primary vector of *S. mansoni* in Lake Victoria, and inhabits the shallow waters and marsh-like habitats of the African Great Lakes and White Nile Watershed (Brown 1994). It has been proposed that *B. sudanica* is conspecific with another species, *B. choanomphala*, which inhabits the deepwater, benthic region of Lake Victoria (Standley et al. 2014; Andrus et al. 2023). However, because the two forms differ widely in their susceptibility to schistosomes (Mutuku et al. 2021), and optimal taxonomic designation would require collections from the type location of *B. sudanica* in Sudan, we use the traditional name *B. sudanica s*.*l*. to refer specifically to the shallow-water form, which is more resistant to schistosome infection than the deepwater type, *B. choanomphala*.

We exposed two lines of *B. sudanica* that were originally collected from Lake Victoria to a line of *S. mansoni* originally obtained from the same region. The exposures represent a compatible and incompatible combination of snail hosts with the parasite line. We compared expression of the five genes among exposed and control snails and across snail lines at 8, 24, and 72 hours post infection. Because a prior study indicated that diet influenced susceptibility of snails to parasites (Trapp et al. 2024), we also compared gene expression between exposed and control snails raised on either a strict low- or high-nutrient diet to determine if diet influenced expression of these immune genes.

We hypothesized that the resistant snail line of *B. sudanica* would have higher constitutive expression of immune genes than susceptible lines or that exposure to parasites would induce higher expression of immune genes compared to susceptible snails, analogous to the work that has been done previously with susceptible (M-line) and resistant (BS-90) lines of *B. glabrata* (Bender et al. 2007; Pila et al. 2016b; Marquez et al. 2022).

## Materials and Methods

### Snails and parasites

This study included two lines of *Biomphalaria sudanica* snails that originate from the Kisumu region of Lake Victoria. All snails were maintained in the laboratory in 10 L plastic shoe boxes with 8 L of aerated and filtered artificial spring water and supplemented with calcium carbonate. Room temperatures were maintained at around 24–25ºC and lights were kept on a 12:12 light-dark cycle. One line, referred to as 163, was purposely inbred after collection from the wild with three generations of selfing. Line 163 is primarily fed commercial snail food pellets (Aquatic Freshwater Snail Food Mix #2 (Aquaticblendedfoods.com)) and supplemented with lettuce. The second line used, KEMRIwu, was not purposely inbred but had been maintained in captivity since 2010. All KEMRIwu snails were fed green leaf lettuce twice a week and supplemented once a week with the commercial food pellets. For the diet comparison study component, we reared KEMRIwu snails on either a strict diet of green leaf lettuce or commercial food pellets (Aquatic Freshwater Snail Food Mix #2 (Aquaticblendedfoods.com)) for at least two generations (as described in (Trapp et al. 2024)).

We used a single line of *S. mansoni* in this study, UNMKenya, that originated from the Lake Victoria region of Kenya. The UNMKenya line of *S. mansoni* shows compatibility with KEMRIwu snails (∼40-50% of exposed snails become infected), whereas line 163 snails are incompatible (<2%) (Spaan et al. 2023). These lines are referred to as susceptible (KEMRI) and resistant (163) throughout the manuscript for readability.

### Snail exposures

All *B. sudanica* snails used in this study were between 5-6mm in shell diameter at the time of *S. mansoni* exposure. Snails were individually exposed to eight freshly hatched miracidia in 2 mL of artificial spring water within the wells of a 24-well tissue culture plate for 3 hours. After exposure, snails were moved to their aquaria and provided food (green leaf lettuce or pellets). Sham-exposed control snails underwent the same procedure but were placed in wells without miracidia and removed after three hours. In total, 108 snails were exposed/sham-exposed to *S. mansoni* in the line comparison study, and 96 in the diet comparison study.

### Identifying immune gene orthologs and qPCR primer design

Primers were designed for the orthologous actin (control/housekeeping gene), biomphalysin, GRN, SOD1, TEP1 and TLR genes (Table 1) using the published *B. sudanica* annotated genome (Pennance et al. 2024). The coding sequences (CDS) for each gene were used to design primers targeting 84-150 bp regions using Primer3 (Untergasser et al. 2012). Primer specificity was determined by searching against the *B. sudanica* genome and CDS sequences for matches using BLAT (Kent 2002) options stepSize=1 minScore=15, that may generate other viable amplicons, and against an alignment of five *B. sudanica* inbred line genomes (Pennance et al. 2024) to infer conserved primer binding regions. An effort was made to design primers for *FREP*2, *FREP*3, and *FREP*4; however, specific primers could not be generated for *FREP*3, and *FREP*4; and those generated for *FREP*2 did not perform adequately, thus these targets were excluded.

**Table 1.**
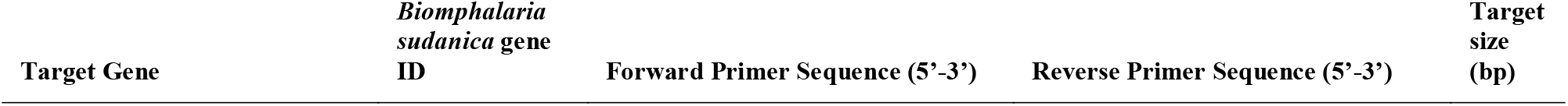
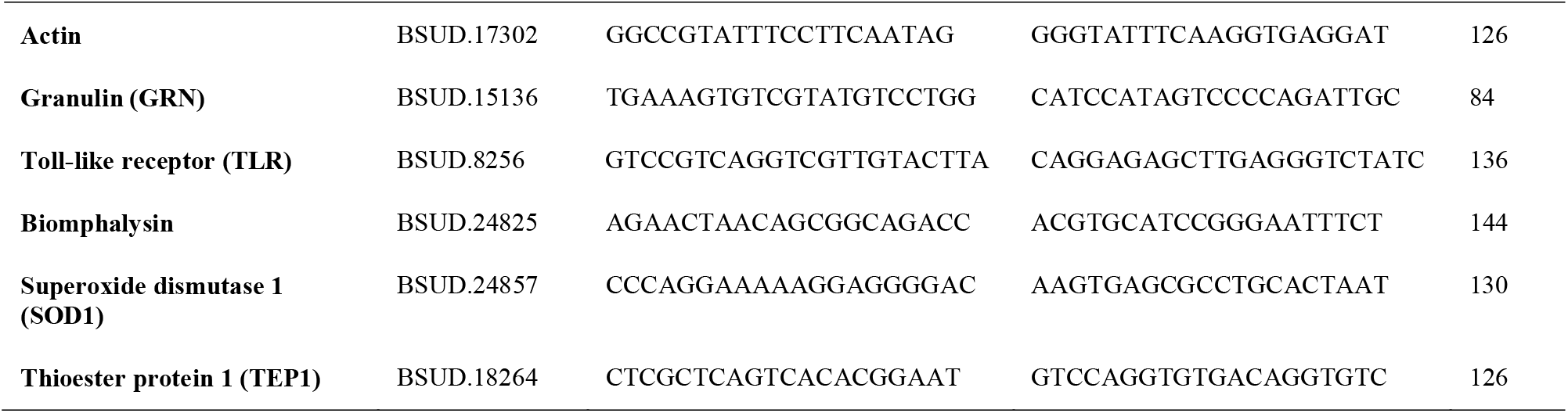
*Biomphalaria sudanica s*.*l*. genes targeted using reverse transcription (RT) quantitative PCR in the current study as a control/housekeeping gene (actin) and five identified as orthologous to anti-schistosome defense genes established in the immune response of *Biomphalaria glabrata* to *Schistosoma mansoni*.

### RNA preservation and extraction

All surfaces were treated with RNase^TM^ Away or RNaseZap^TM^ RNase Decontamination Reagent/Solution/Wipes (Invitrogen, MA, USA), prior to snail sacrifice and preservation. At the designated time points post-parasite exposure, snails were individually removed from their respective aquaria and placed directly in RNA/DNA free Omni 2mL Reinforced Tubes containing 2.8mm ceramic beads (Omni International, Kennesaw GA, US) and 0.5 mL of TRIzol^TM^ reagent (Invitrogen, MA, USA). The sealed tube was immediately placed in an Omni Beadrupter 12 (Omni International, Kennesaw GA, US) and tissue homogenized (speed = 6.00, cycle time = 5 seconds, number of cycles = 1). The homogenized samples were then stored at - 80ºC.

For RNA extraction, frozen homogenized samples were thawed on wet ice and then incubated at room temperature for 5 minutes before proceeding with a modified Trizol PureLink^TM^ (Invitrogen, MA, USA) RNA extraction protocol (detailed in full in Supplement 1). In brief, phase separation was performed with chloroform:isoamyl alcohol retaining the RNA aqueous phase that was then bound to a PureLink column membrane. A PureLink DNase (Invitrogen, MA, USA) on-column treatment was performed, RNA washed, and purified RNA eluted from membrane in RNA/DNA/RNase free water. If qPCR was to be conducted the same day, the extracted RNA was reverse transcribed (see below) immediately following 5 minutes of incubation at room temperature. Otherwise the extracted RNA samples were stored at -80 C.

### Reverse transcription quantitative PCR (RT-qPCR) of candidate immune genes

Total purified RNA was quantified using Qubit RNA Broad Range (BR) Assay Kit using the Qubit 4 Flourometer (Invitrogen, MA USA). Purity and presence of contaminants in RNA samples were assessed by measuring 260/230 and 260/280 absorbance values on a Nanodrop to check that samples were within the expected range for pure RNA, ∼1.8-2.2 and ∼2.0, respectively.

Reverse transcription of up to a total of 1 µg of RNA per sample per reaction was performed using qScript cDNA SuperMix (Quantabio, MA, USA) following the manufacturers protocol. In brief, per 0.2 ml reaction tube, 4 µl of qScript cDNA SuperMix were added with a volume of total RNA up to a total mass of 1 µg and the remaining volume to make up to 20 µl was made up of RNA/DNA free water. For samples with an RNA concentration <62.5 ng/µl, 16 µl of RNA extract were used without the addition of water. Cycling parameters for reverse transcription followed those recommended in the manufacturers protocol (5 mins at 25 C, 30 mins at 42 C, 5 minutes 85 C, hold at 4 C). cDNA was used immediately following reverse transcription, or stored at -20 C for up to 1 week.

qPCRs were performed in 20 µl reactions in 0.1 mL MicroAmp^TM^ Optical 96-well reaction plates (Applied Biosystems, MA, USA). Each reaction contained 10 µl of PowerUp^TM^ SYBR^TM^ Green Master Mix (Applied Biosystems, MA, USA), 0.5 µl of each of the respective forward and reverse primer (10 µM, diluted in TE buffer) and 8 µl of RNA/DNA free water. For each qPCR plate set up and run, samples were chosen to represent each experimental group within the respective study. Each sample was run with the housekeeping gene (actin) and up three of the five target genes at a time (either GRN, TLR or biomphalysin, SOD1, TEP1). Each sample was run in triplicate for each target gene on the same plate. Therefore, per plate, each sample occupied 12 wells per plate (4 targets, 3 technical replicates), with a total of 8 samples per 96-well plate. Gene expression for each reaction (including qPCR efficiency, see *Determining primer qPCR efficiency* below) was inferred by setting the delta Rn (relative sample fluorescence) threshold to 0.2 across all experiments, since this represented a reliable exponential portion of the fluorescence curve. From the 0.2 delta Rn threshold, the cycle threshold (CT) data was measured for each sample. The CT values were averaged across the technical replicates to reduce technical errors, excluding significant outlier technical replicates that could be due to significant technical errors (e.g. pipetting error) (See Supplement 2). As a result, in some instances, CT values were averaged from two replicates instead of three when one replicate was identified as an outlier (detailed in datasets: Supplementary Table 2 and 3). From the average CT values across replicates, delta CT (ΔCT) was calculated as the difference between the CT of each target gene and the CT of the housekeeping gene (actin) for each snail, providing a measure of relative gene expression.

### Determining primer qPCR efficiency

The efficiency of qPCR primers was evaluated to ensure they fell within the optimal range (90-110%) for each target gene. To do this, cDNA from a KEMRIwu snail used in the study (study ID: KK8_6) was generated from 1 µg of RNA and diluted by a factor of 1:5 three times to generate cDNA template concentrations of full, 1:5, 1:25 and 1:125. The qPCR was performed for actin and each target gene (Table 1) using each diluted template in triplicates. The triplicate CT values were averaged for each gene, and then the average CT was plotted against the log of sample quantity to calculate the slope of the regression. The efficiency percentage per target was then calculated using the efficiency percentage equation: Efficiency percentage = (10^-1/regression^-1)x100. All qPCR efficiencies were between 94-100% (Supplementary Table 1).

### Statistical analysis

All data handling and statistical analyses were performed in R v4.4.1, with statistical significance determined as *p* < 0.05. Prior to comparative analysis, major ΔCT outliers were removed from each dataset. Major outliers were defined as any values falling below 1.5 times the interquartile range (IQR) (below the 25th percentile) or above 1.5 times the IQR (the 75th percentile) for each snail line or diet group, per target gene, at each time point. Data normality for each target gene and treatment group was assessed using the Shapiro-Wilk test and by the examining residual plots from a fitted linear model (LM) with an interaction between the snail exposure to *S. mansoni* (Condition) and the snail line or diet group (lm(deltaCT∼Condition*SnailLine/Diet). Data was considered normal if the Shapiro-Wilk *p* value was non-significant (*p* ≥ 0.05), but if significant, the residual plots and histograms from the LM were used to infer normality. Following these criteria, all data were normally distributed.

A generalized linear model (GLM) of Gaussian family was fitted to the two datasets using an interaction between condition and snail line / diet group. Models with significant interaction terms were run a second time changing the reference level to interpret the direction of the interaction. All models with non-significant interaction terms were rerun using variables as fixed effects only, and this result was then used to infer gene expression differences between groups.

## Results

### Comparison of induced and constitutive gene expression in susceptible vs. resistant snails

In total, 578 ΔCT values from between 3 and 12 individual snails per gene and experimental group, were used for the analysis of candidate gene expression between susceptible and resistant snails (Table 3).

**Table 3.**
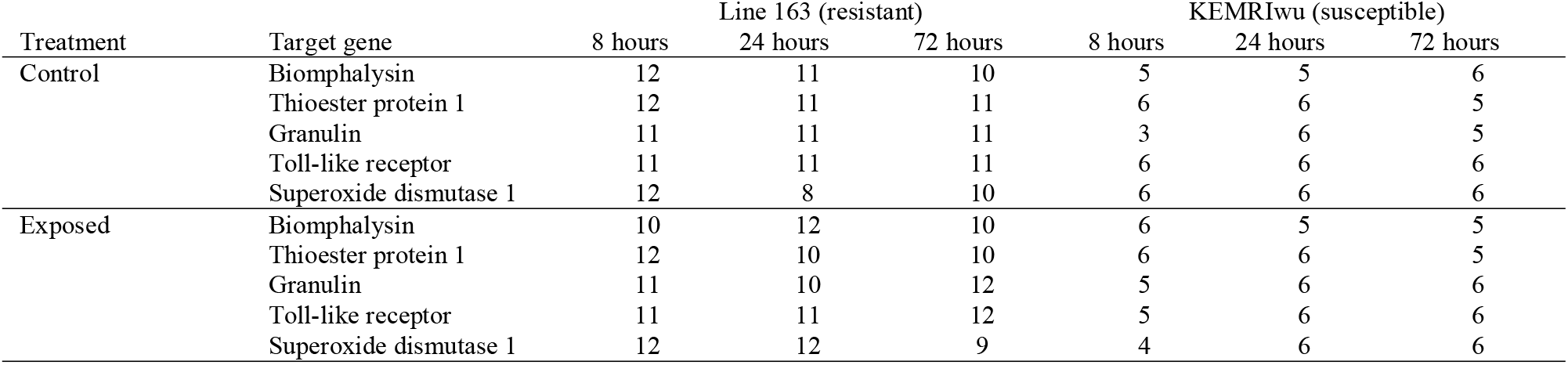
Sample sizes per target gene for *Biomphalaria sudanica* line 163 and KEMRIwu snails sham exposed (Control) or exposed to *Schistosoma mansoni* (Exposed) in the snail line comparative study.

Eight hours post experimental treatment, where snails were either exposed or sham-exposed to *S. mansoni*, both susceptible (KEMRIwu) and resistant (Line 163) *B. sudanica* had significantly higher expression of GRN (Figure 1c) and SOD1 (Figure 1e) induced by *S. mansoni* exposure (GLMs FE models, Table 4). Also at 8 hours post exposure, significant interaction terms in GLMs for biomphalysin (GLM, β = -1.9094, *p* = 0.0047, Supplementary Table 4) and TEP1 (GLM, β = -1.5529, *p* = 0.0010, Supplementary Table 4) demonstrated that for both genes significant upregulation occurred in susceptible snails exposed to *S. mansoni*, whereas no significant change was observed in resistant snails when exposed to *S. mansoni* (Figure 1a and 1b). In addition, exposed susceptible snails had a significantly higher expression of both biomphalysin and TEP1 than exposed resistant snails (Figure 1a and 1b).

**Table 4.** Results of generalized linear models (GLM) performed with interactions (INT), and fixed effects (FE) for INT models with a non-significant interaction term, for each of the five *Biomphalaria sudanica* candidate immune genes for *Schistosoma mansoni* resistance. GLMs assess the constitutive and induced gene expression differences in *B. sudanica* that are either resistant (Line 163) or susceptible (KEMRIwu) to *S. mansoni*. The reference levels are the same for each INT and FE model, schistosome exposure = sham-exposed (control) and snail line = resistant (Line 163), with results for INT models with alternative references given in Supplementary Table 4. In GLMs, positive estimate values for schistosome exposure represent a decrease in gene expression (i.e. increased ΔCT) in exposed snails compared to the reference sham-exposed snails, whereas positive estimate values for snail line represent a lower gene expression in susceptible snails than resistant snails.

| Gene | GLM | Predictors | 8 hours |  | 24 hours |  | 72 hours |  |
| --- | --- | --- | --- | --- | --- | --- | --- | --- |
| | | | Estimate ± SE | $p^1$ | Estimate ± SE | $p^1$ | Estimate ± SE | $p^1$ |
| Biophalysin | INT | Intercept | 1.94 ± 0.24 | *** | 2.97 ± 0.24 | *** | 2.14 ± 0.19 | *** |
| | | Schistosome exposure (exposed) | 0.57 ± 0.36 | | -0.76 ± 0.34 | * | 0.40 ± 0.27 | |
| | | Snail line (susceptible) | 0.86 ± 0.45 | | -0.75 ± 0.44 | | -0.35 ± 0.31 | |
| | | Schistosome exposure – snail line | -1.91 ± 0.62 | ** | 1.37 ± 0.61 | * | 0.29 ± 0.45 | |
| | FE | Intercept | | | | | 2.09 ± 0.17 | *** |
| | | Schistosome exposure (exposed) | | | | | 0.50 ± 0.21 | * |
| | | Snail line (susceptible) | | | | | -0.21 ± 0.22 | |
| Thioester protein 1 (TEP1) | INT | Intercept | 5.86 ± 0.23 | *** | 5.74 ± 0.22 | *** | 4.84 ± 0.34 | *** |
| | | Schistosome exposure (exposed) | -0.34 ± 0.33 | | 0.28 ± 0.32 | | -0.18 ± 0.49 | |
| | | Snail line (susceptible) | 0.19 ± 0.40 | | 0.02 ± 0.37 | | 0.52 ± 0.61 | |
| | | Schistosome exposure – snail line | -1.55 ± 0.57 | ** | -0.36 ± 0.52 | | -0.41 ± 0.86 | |
| | FE | Intercept | | | 5.80 ± 0.20 | *** | 4.90 ± 0.31 | *** |
| | | Schistosome exposure (exposed) | | | 0.15 ± 0.25 | | -0.32 ± 0.40 | |
| | | Snail line (susceptible) | | | -0.16 ± 0.26 | | 0.32 ± 0.43 | |
| Granulin (GRN) | INT | Intercept | 4.36 ± 0.32 | *** | 3.98 ± 0.25 | *** | 5.94 ± 0.55 | *** |
| | | Schistosome exposure (exposed) | -0.71 ± 0.45 | | 0.05 ± 0.36 | | -0.67 ± 0.76 | |
| | | Snail line (susceptible) | 0.69 ± 0.68 | | 0.81 ± 0.41 | | -0.86 ± 0.98 | |
| | | Schistosome exposure – snail line | -0.40 ± 0.89 | | -0.12 ± 0.59 | | -0.10 ± 1.33 | |
| | FE | Intercept | 4.41 ± 0.29 | *** | 4.00 ± 0.22 | *** | 5.96 ± 0.49 | *** |
| | | Schistosome exposure (exposed) | -0.81 ± 0.38 | * | 0.01 ± 0.28 | | -0.70 ± 0.61 | |
| | | Snail line (susceptible) | 0.45 ± 0.43 | | 0.76 ± 0.29 | * | -0.92 ± 0.65 | |
| Toll-like receptor (TLR) | INT | Intercept | 7.96 ± 0.47 | *** | 8.00 ± 0.39 | *** | 9.82 ± 0.63 | *** |
| | | Schistosome exposure (exposed) | -0.60 ± 0.67 | | -0.23 ± 0.55 | | -1.10 ± 0.87 | |
| | | Snail line (susceptible) | 0.79 ± 0.79 | | 1.43 ± 0.65 | * | 0.59 ± 1.05 | |
| | | Schistosome exposure – snail line | -1.35 ± 1.16 | | 0.19 ± 0.93 | | -0.15 ± 1.48 | |
| | FE | Intercept | 8.18 ± 0.43 | *** | 7.97 ± 0.35 | *** | 9.85 ± 0.56 | *** |
| | | Schistosome exposure (exposed) | -1.05 ± 0.55 | | -0.17 ± 0.44 | | -1.16 ± 0.69 | |
| | | Snail line (susceptible) | 0.16 ± 0.58 | | 1.52 ± 0.46 | ** | 0.51 ± 0.73 | |
| Superoxide dismutase (SOD1) | INT | Intercept | 1.57 ± 0.08 | *** | 1.43 ± 0.12 | *** | 1.38 ± 0.08 | *** |
| | | Schistosome exposure (exposed) | -0.10 ± 0.12 | | 0.03 ± 0.16 | | -0.02 ± 0.12 | |
| | | Snail line (susceptible) | 0.35 ± 0.14 | * | 0.38 ± 0.19 | | 0.33 ± 0.13 | * |
| | | Schistosome exposure – snail line | -0.42 ± 0.22 | | -0.08 ± 0.26 | | 0.15 ± 0.19 | |
| | FE | Intercept | 1.63 ± 0.08 | *** | 1.45 ± 0.11 | *** | 1.35 ± 0.07 | *** |
| | | Schistosome exposure (exposed) | -0.22 ± 0.10 | * | 0.00 ± 0.12 | | 0.04 ± 0.09 | |
| | | Snail line (susceptible) | 0.17 ± 0.11 | | 0.34 ± 0.13 | * | 0.40 ± 0.10 | *** |

<sup>1</sup> $p$ values represent: \*\*\* = <0.001, \*\* = <0.01, \* = <0.05.

**Figure 1.**
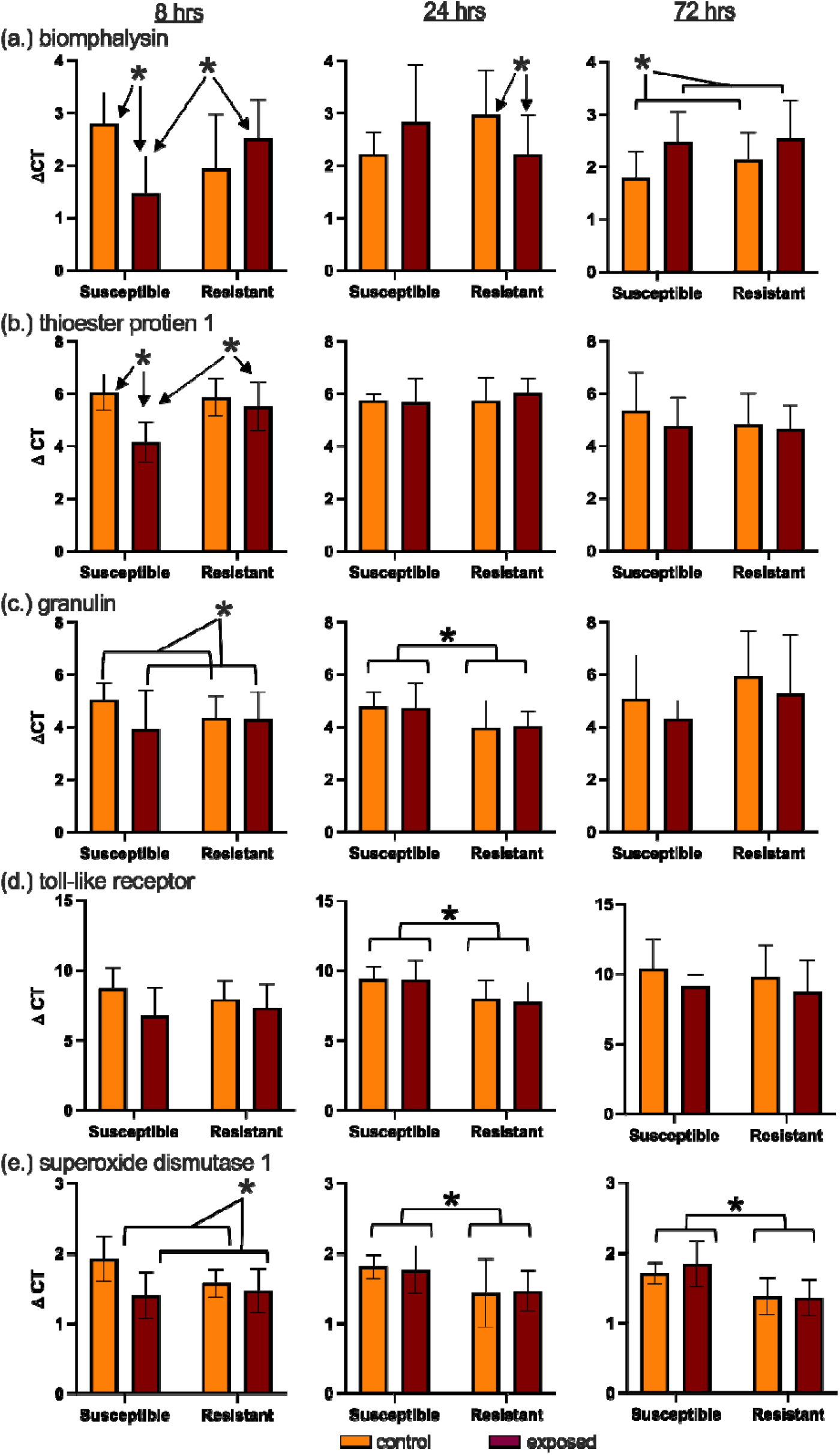
Expression of 5 candidate resistance genes of KEMRIwu (susceptible) and 163 (resistant) line of *Biomphalaria sudanica* that were either challenged with *Schistosoma mansoni* (exposed) or sham exposed (control). Expression is represented by delta cycle threshold (ΔCT), and is relative to the expression of actin. Note that gene expression and ΔCT have an inverse relationship. Expression was measured at 8, 24, and 72 hours for: a) biomphalysin, b) thioester protein 1, c) granulin, d) toll-like receptor, and e) superoxide dismutase 1. Mean ΔCT and standard deviation for snails in each experimental group are shown, and asterisks indicate a statistically significant difference, with square brackets representing significance from FE models and arrows indicate interaction effects (see Table 4).

Following 24 hours post-exposure to *S. mansoni*, a significant interaction was observed in the GLM for biomphalysin, which upon further interrogation represents a significant upregulation of biomphalysin in exposed resistant snails compared to control resistant snails (Figure 1a, Supplementary Table 4). Furthermore, resistant snails had relatively higher constitutive expression of genes GRN (Figure 1c), SOD1 (Figure 1e), and TLR (Figure 1d) at 24 hours compared to susceptible snails and independent of *S. mansoni* exposure (Table 4). The constitutively higher expression of SOD1 in resistant snails also persisted at 72 hours (Figure 1e), at which point the only parasite induced gene expression change was in biomphalysin, that showed significantly lower expression in exposed snails compared to controls (Figure 1a).

### Comparison of gene expression in snails on different diets

In total, 256 ΔCT values obtained from between 8 and 12 snails per experimental group were used for the analysis of induced and constitutive gene expression of three candidate genes from lettuce fed and pellet fed KEMRIwu snails either exposed to *S. mansoni* or control (Table 4).

**Table 4.** Number of delta CT (ΔCT) values from individual *Biomphalaria sudanica* KEMRIwu (susceptible) snails used per target gene per experimental group in the diet comparative study.

| Experimental group | Gene | KEMRIwu lettuce fed |  | KEMRIwu pellet fed |  |
| --- | --- | --- | --- | --- | --- |
|  |  | 8 hours | 24 hours | 8 hours | 24 hours |
| Control | Biomphalysin | 11 | 11 | 12 | 11 |
|  | Superoxide dismutase 1 | 12 | 11 | 11 | 11 |
|  | Thioester protein 1 | 11 | 11 | 11 | 10 |
| Exposed | Biomphalysin | 11 | 9 | 10 | 11 |
|  | Superoxide dismutase 1 | 11 | 10 | 11 | 11 |
|  | Thioester protein 1 | 8 | 10 | 10 | 11 |

Diet induced gene expression was measured for genes SOD1, biomphalysin, and TEP1 at 8 and 24 hours post-exposure to *S. mansoni* or sham-exposure. The hypothesis tested was that gene expression, either induced or constitutive, would be greater in snails fed a lettuce diet compared to those fed a pellet diet, because a previous study (Trapp et al. 2024) showed that lettuce fed snails were more resistant to infection. Expression of SOD1 was consistent with this hypothesis, with lettuce-fed snails showing higher constitutive expression compared to pellet snails at both 8 and 24 hours (Figure 2, Table 5). Schistosome exposure did not induce expression differences (Table 5).

**Table 5.**
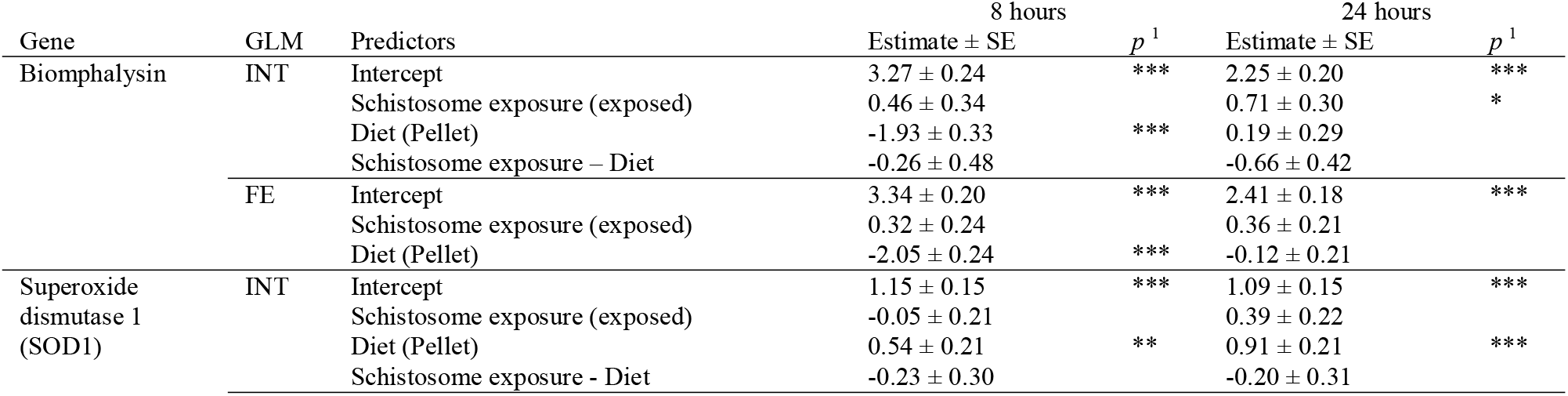
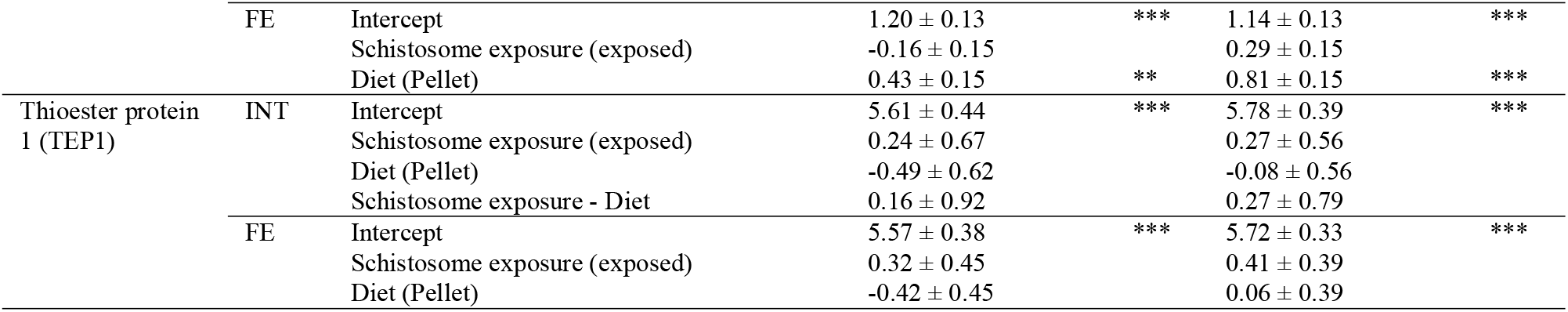
Results of generalized linear models (GLM) performed with interactions (INT), and fixed effects (FE) for INT models with a non-significant interaction term, for three *Biomphalaria sudanica* candidate immune genes for *Schistosoma mansoni* resistance. GLMs assess the constitutive and induced (via schistosome exposure) gene expression differences in susceptible *B. sudanica* KEMRIwu inbred line snails that are fed solely a diet of green leaf lettuce or commercial pellets. In GLMs, positive estimate values for schistosome exposure represent a decrease in gene expression (i.e. increased ΔCT) in exposed snails compared to the reference control snails, whereas positive estimate values for diet represent a lower gene expression in pellet fed snails than lettuce fed snails.

**Figure 2.**
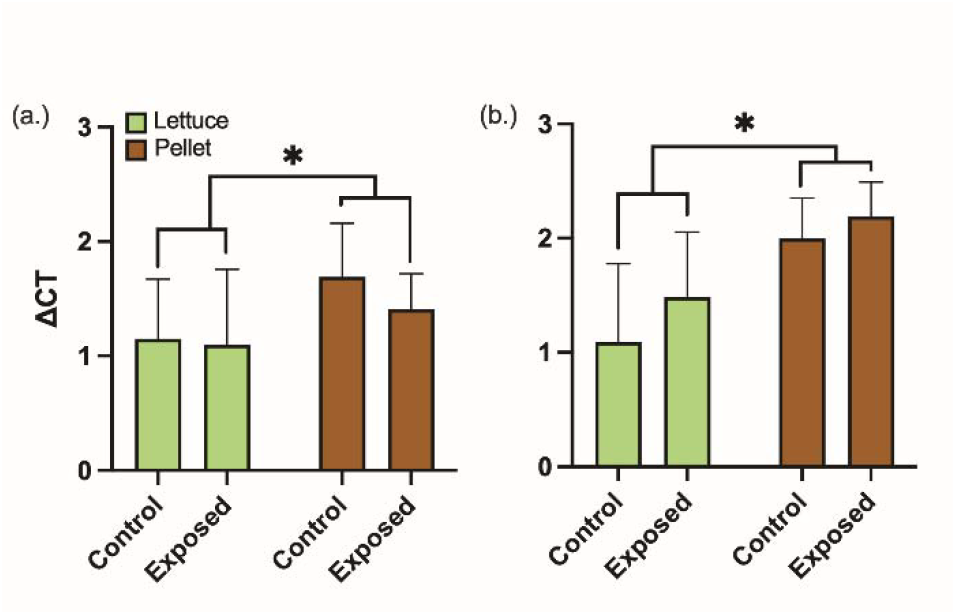
Relative expression levels of superoxide dismutase 1 (SOD1) gene of *Biomphalaria sudanica* KEMRIwu line fed exclusively lettuce or pellet food that were either challenged with *Schistosoma mansoni* (Exposed) or sham-exposed (Control). Delta cycle threshold (ΔCT) is relative to the expression of the housekeeping gene, actin. Expression was measured at a) 8 hours and b) 24 hours post-exposure. Mean ΔCT and standard deviation are shown, and asterisks indicate a statistically significant difference (p<0.05).

At 8 hours, biomphalysin expression in pellet fed snails was constitutively higher than that of lettuce fed snails; however, no expression differences in biomphalysin were detected at 24 hours and no induced expression differences from schistosome exposure were detected in biomphalysin (Table 5), despite our previous findings from KEMRIwu (susceptible) snails fed a mixed diet (Figure 1a). No induced or constitutive expression differences were detected in TEP1 despite our previous findings of induced changes at 8 hours (Figure 1b).

## Discussion

With the goal of characterizing the anti-schistosome defense of an African snail vector of schistosomiasis, we measured expression of five putative immune genes in two genetic lines of *B. sudanica* that are compatible or incompatible with *S. mansoni*. Expression was measured at three time points post schistosome exposure and compared to sham-exposed controls. Furthermore, we characterized how diet impacts gene expression since a pellet diet was shown to increase susceptibility of *B. sudanica* compared to those fed a lettuce diet (Trapp et al. 2024). Overall, we found that our results were only partially concordant with expectations based on previous results of the *B. glabrata-S. mansoni* system.

A role for SOD1 in anti-schistosome defense was strongly supported by the expression data. The resistant snail line showed higher expression than the susceptible snails at the 24 and 72hour time points regardless of exposure to *S. mansoni*. Additionally, induced expression of SOD1 was detected at 8 hours post-exposure, with both susceptible and resistant exposed snails showing greater expression than control snails. The results of the diet experiment further corroborated the role of SOD1 in anti-schistosome defense in that those fed the pellet diet, which yields more susceptible snails (Trapp et al. 2024), showed lower expression of SOD1 than those fed the lettuce diet, which yields more resistant snails. Surprisingly, induced expression was not observed in the diet dataset. This may suggest a brief window of SOD1 induction following schistosome exposure that was not captured in the diet study, or a potential impact of dietary differences between the two studies. Previous work has linked greater expression of SOD1 to schistosome resistance in *B. glabrata* (Goodall et al. 2004, 2006; Bender et al. 2007). SOD1 is expressed by hemocytes and catalyzes the conversion of superoxide anion to H_2_O_2_, which is involved in killing of parasite sporocysts in resistant *B. glabrata* (Hahn et al. 2001a). While these findings support that SOD1 expression is involved in the anti-schistosome defense of *B. sudanica*, it clearly is not the only factor that renders snails resistant, as here we show evidence that schistosome exposure induced similar levels of SOD1 in both susceptible and resistant snails.

Granulin expression was induced in both susceptible and resistant *B. sudanica* at 8 hours post-exposure. Early induced expression of granulin is consistent with the role of this growth factor in stimulating proliferation and differentiation of hemocytes for pathogen defense, as demonstrated further in siRNA mediated knockdown studies of GRN in *B. glabrata* that can result in a loss of resistance (Hambrook et al. 2020). In previous work with *B. glabrata*, after exposure to *S. mansoni*, granulin expression was significantly different in exposed resistant snails (BS-90) compared to matched controls 12-48 hours post exposure, and at 24 hours in exposed susceptible snails (M-line) compared to controls (Pila et al. 2016a). Thus, the results are concordant in that there is an early peak of granulin expression in susceptible and resistant snails post-exposure, however, the peak appears earlier for *B. sudanica*.

TEP1, a putative opsonin that is highly expressed in snail hemocytes that binds to *S. mansoni* sporocysts and promotes phagocytosis of the pathogen (Li et al. 2020), was only seen to increase in expression in the susceptible snails 8 hours after exposure in this study. A previous study of *B. glabrara* (albino Brazilian snails) showed upregulation 6-24 hours post-exposure to incompatible schistosomes (Portet et al. 2018). These results were not replicated in a different study using M-line and BS90 *B. glabrata* and, in fact, the susceptible snail (M-line) showed higher constitutive expression of TEP1 than the resistant snail (BS90) (Marquez et al. 2022). However, in this same study, several TEP paralogs were differentially regulated post-exposure, and therefore, these paralogs may swap roles with TEP1 in anti-schistosome defense of different lines of *B. glabrata* (Marquez et al. 2022). Future work should investigate the roles of the TEP paralogs in anti-schistosome defense of *B. sudanica*, and determine whether they play a similar role in forming a functional complex with FREP3 proteins to promote production of reactive oxygen species (Li et al. 2020).

Biomphalysin-1 showed induced expression in susceptible *B. sudanica* at 8 hours, induced expression in resistant snails at 24 hours, and constitutive differences at 72 hours in that susceptible snails had higher expression than resistant snails. These data indicate that this biomphalysin is induced by schistosome exposure in *B. sudanica*, and may be differentially expressed among susceptible and resistant lines; however, higher expression is not correlated with resistance in this case. Biomphalysin-1 has the structure of an aerolysin pore forming toxin and has been shown to bind to the surface of *S. mansoni* sporocysts (Galinier et al. 2013). Additionally, it forms a complex with the aforementioned TEP1 and other fibrinogen related proteins in resistant snails (Li et al. 2020). Previous studies have indicated that expression of biomphalysin-1 is not induced by schistosome exposure (Galinier et al. 2013); however, other biomphalysin paralogs are induced upon exposure to schistosomes (4 and 21) and other pathogens (Pinaud et al. 2021). Further work on this family of pore forming toxins will help unravel which may be specific to the anti-schistosome defense of *B. sudanica*.

In this study, TLR showed no induced expression upon challenge with *S. mansoni*. There was evidence that resistant snails showed higher constitutive expression; however, this observation only occurred at 24 hours. Previous work has shown mixed findings in *B. glabrata*, including higher constitutive TLR expression in resistant snails compared to susceptible snails, as well as a strong induction of TLR expression in resistant snails and a more modest response in susceptible snails following schistosome challenge (Hanington et al. 2010; Pila et al. 2016a; Zhong et al. 2024). Given the strength of previous studies, it is surprising that induced expression of TLR was not detected in *B. sudanica* snails after schistosome challenge. There are multiple copies of toll-like receptors in *B. sudanica* and it is possible that the function in the anti-schistosome response has been shifted to a paralog TLR.

Overall, despite the vast evidence that these five genes (Biomphalysin, GRN, SOD1, TEP1 and TLR) have a strong role in the immune defense of *B. glabrata* against *S. mansoni*, the presented expression data do not present a clear role for all of them in the anti-schistosome defense of *B. sudanica*. African *Biomphalaria* diverged from a common ancestor with *B. glabrata* less than 5 million years ago (Bu et al. 2023), and thus conservation of their immune response might be expected. However, in this short time, African *Biomphalaria* diverged into ∼12 species across the continent and were exposed to a vast number of novel pathogens including *S. mansoni*, which is highly diverse in East Africa (Platt et al. 2022). At the same time,

*S. mansoni* is a relatively new pathogen to *B. glabrata* as it was transported from Africa to South America and the Caribbean with the 16^th^-19^th^ Century Atlantic Slave Trade (Fletcher et al. 1981; Després et al. 1993; Crellen et al. 2016). Thus, local adaptation and coevolution of hosts and pathogens could have driven differences in the basic anti-schistosome response of these snails. Many of these candidate immune genes belong to diversified families that likely arose through gene duplication. It is possible that the role of some of these immune genes has shifted to these paralogs, or perhaps to other genes yet unrecognized.

Another possibility is that expression variation in the susceptible line could have masked some induced expression patterns. Unlike the *B. glabrata* system where susceptible snails exhibit nearly 100% susceptibility, the KEMRIwu *B. sudanica* is only partially susceptible with a dose of 5 miracidia yielding a prevalence of 35-50% (Spaan et al. 2023; Trapp et al. 2024).

In conclusion, while these findings support a conserved role of some of the characterized anti-schistosome immune genes of *B. glabrata*, they do not provide strong evidence for the involvement of each gene. Given that *S. mansoni* is vectored by multiple *Biomphalaria* species, a comprehensive understanding of host-vector compatibility requires investigation of non-model vectors such as *B. sudanica*. Expanding genomic and transcriptomic resources for these species is essential to facilitate further research in this area. (Pennance and Rollinson 2024). These resources would enable robust functional transcriptomic studies to assess the expression of multiple genes and identify a broader set of genes potentially responsible for schistosome resistance. This could pave the way for targeted gene silencing studies, leveraging techniques that have already been established in *B. glabrata* inbred lines.

## Supporting information

Supplement 1 RNA extraction

Supplement 2 Outlier removal

## Acknowledgements

Stephanie Bollmann (Oregon State University) for the initial design of primers TLR, GRN and Actin.

## Financial support

This work was supported by the National Institutes of Health, Grant/Award Number: R01AI141862.

## Conflict of interest declaration

The authors declare no conflicts of interest.

## Ethics statement

Project approval received under the relevant bodies, including KEMRI Scientific Review Unit (permit # KEMRI/RES/7/3/1), Kenya’s National Commission for Science, Technology, and Innovation (permit # NACOSTI/P/22/148/39 and NACOSTI/P/15/9609/4270), Kenya Wild-life Services (permit # WRTI-0136-02-22 and # 0004754), and National Environment, Management Authority (permit # NEMA/AGR/159/2022 and # NEMA/AGR/46/2014) (Registration # 0178). The UNMKenya parasite strain was maintained in hamsters at the Western University of Health Sciences under approval of the IACUC (ACUP# R20IACUC039).

## Supplementary material

**Supplement 1**. Protocol for the modified Trizol PureLink^TM^ (Invitrogen, MA, USA) RNA extraction protocol used to extract all RNA in the current study.

**Supplement 2**. Detailed methods of removal of significant CT outliers within technical replicates and delta cycle threshold (ΔCT) outliers.

**Supplementary Table 1**. Results of qPCR primer (see Table 1) efficiency tests for each target gene.

**Supplementary Table 2**. Delta cycle threshold (ΔCT) results of snail line comparative gene expression analysis with outliers marked.

**Supplementary Table 3**. Delta cycle threshold (ΔCT) results of diet comparative gene expression analysis with outliers marked.

**Supplementary Table 4**. Results of generalized linear models with significant interaction terms in snail line comparative gene expression analysis with changing reference levels.

## Notes

### Competing Interest Statement

The authors have declared no competing interest.

