## Supplement 1 RNA extraction for "Schistosome exposure and diet induced effects on candidate immune gene expression in an African snail vector"

**Supplement 1. Modified Trizol^TM^ PureLink RNA extraction protocol for use with *Biomphalaria* snails**

**Procedural guidelines**

- Perform all steps at room temperature (20–25°C) unless otherwise noted.
- Use disposable, individually wrapped, sterile plastic ware and sterile, disposable RNase-free pipettes, pipette tips, and tubes.
- Wear disposable gloves while handling reagents and RNA samples to prevent RNase contamination from the surface of the skin; change gloves frequently, particularly as the protocol progresses from crude extracts to more purified materials.
- Always use proper microbiological aseptic techniques when working with RNA.
- Use RNase*Zap*TM RNase Decontamination Solution (Cat. no. AM9780) to remove RNase contamination from work surfaces and non-disposable items such as centrifuges and pipettes used during purification.

**1. Lyse samples**

a. Set refrigerated centrifuge to 4C.
b. Add 0.5 mL of TRIzolTM Reagent to RNA/DNA free sterile OMNI bead rupting tube with 2.8 mm ceramic beads for hard tissue homogenization (10032-756). Add 4-5 mm snail (contains ~20 mg of tissue) and homogenize in the bead ruptor at a speed of 6.00 for 5 seconds. Store immediately in -80C freezer or continue to step b.
c. Incubate homogenized sample for ~5 minutes (~10 minutes if from frozen) to permit complete dissociation of the nucleoproteins complex.
d. Add 0.2 mL of chloroform:isoamyl alcohol (24:1), then securely cap the tube and gently invert to mix.
e. Incubate (room temp) for 2–3 minutes.
f. Centrifuge the sample for 15 minutes at 12,000 × *g* at 4°C. (The mixture separates into a lower red phenol-chloroform, and interphase, and a colorless upper aqueous phase)
g. Carefully remove tube from centrifuge keeping at angle. Transfer ~330 μL of the colorless, upper aqueous phase containing the RNA to a new tube. Do not disturb other layers!
h. Add an equal volume (~330 µL) of 70% ethanol (room temp), then mix well by gentle vortexing.
i. Invert the tube to disperse any visible precipitate that may form after adding ethanol.

**2. Bind the RNA to the membrane**

Use room temperature centrifuge

a. Transfer up to 700 μL (i.e. 660) of the sample to a spin cartridge (with collection tube)
b. Centrifuge at 12,000 × *g* (room temperature) for 15 seconds.
c. Discard the flow-through, then reinsert the spin cartridge into the same collection tube. (Repeat step 2a–step 2c until the entire sample has been processed if more than 700 µL of starting material).

**3. Washing RNA and DNase treatment**

Before beginning, prepare PureLink® DNase for on-column treatment, add the following components (supplied with PureLink® DNase Set 12185010) to a clean, RNase-free microcentrifuge tube. Prepare 80 μL of DNase reaction per sample in an RNA/DNA/RNase-free microcentrifuge tube:

**10X Dnase I Reaction Buffer	8 µL	[room temp]**

**Dnase (~3U/μL)	10 µL	[4C fridge / -20 for stored aliquots]**

**Rnase Free Water	62 µL	[room temp]
Final Volume	80 µL**

a. Add 350 μL Wash Buffer I to the Spin Cartridge containing the bound RNA (see sample–specific protocol). Centrifuge at 12,000 × g for 15 seconds at room temperature. Discard the flow-through and the Collection Tube. Insert the Spin Cartridge into a new Collection Tube.
b. Add 80 μL PureLink® Dnase mixture (prepared as described above in bold) directly onto the surface of the Spin Cartridge membrane.
c. Incubate at room temperature for 15 minutes.
d. Add 350 μL Wash Buffer I to the Spin Cartridge. Centrifuge at 12,000 x g for 15 seconds at room temperature. Discard flow-through and the Collection Tube and insert the Spin Cartridge into a new Collection Tube.
e. Add 500 μL Wash Buffer II (with ethanol added) to the Spin Cartridge.
f. Centrifuge at 12,000 x g for 15 seconds at room temperature. Discard flow-through and reinsert the Spin Cartridge into the same Collection Tube.
g. Repeat Steps e–f, one additional time, for a total of two washes with Wash Buffer II.

**4. Elute the RNA**

a. Centrifuge at 12,000 × *g* for 1 minute to dry the membrane.
b. Discard the collection tube, then insert the spin cartridge into the **FINAL** RNA/DNA/RNase-free microcentrifuge tube.
c. Add 50 μL of RNase-free water to the center of the spin cartridge.
d. Incubate for 1 minute.
e. Centrifuge at >12,000 × *g* for 2 minutes.
f. Discard the spin cartridge.

The recovery tube contains the purified total RNA.

**5. Measure RNA concentration and quality**

Follow QuBit Broad Range RNA assay and nanodrop (for 260/230 [aim for ~2.0 or above] and 260/280 [pure RNA is ~2.0] measurements) guides. Repeat measures (2-3) on QuBit BR assay and then take an average of the measures to calculate the input RNA amount for cDNA generation.

**6. Store RNA**

Store the purified RNA on ice if used within a few hours. For long-term storage, store the purified RNA at –80°C.
