## Supplement 2 Outlier removal for "Schistosome exposure and diet induced effects on candidate immune gene expression in an African snail vector"

**Supplement 2. Removal of significant CT outliers within technical replicates and delta CT outliers**

The results of the quantitative PCR relies on 3 technical replicates to help account for technical errors such as pipetting errors. The CT values of the technical replicates are averaged together for analysis. However, in some cases, one of the individual technical replicates deviated substantially from the other two. Our rationale for using an outlier analysis to identify such values is that the average of the technical replicates are more likely to reflect true biological values if the outliers are excluded. The outlier analysis was based on the distribution of standard deviations of CT among technical replicates. We pooled the data from this study along with a pilot dataset so that the distributions of standard deviation would be robust.

The outlier analysis was based on the distributions of the standard deviations of the CT values for each target gene in a large dataset that included the current data and pilot data so that estimates of the distribution would be robust. A Gaussian model was fit to the frequency data of the standard deviation for samples with a CT standard deviation <1. The standard deviation for the difference between normally distributed data is √2 * σ, with σ (sigma) representing the prominent peak in the Gaussian model, aka the standard deviation of the data. The 95% confidence interval of the difference, calculated as twice the standard deviation (2[√2 * σ]), was used as the threshold for the inclusion/exclusion of CT data within each replicate group for the data in the current study. The Gaussian fitting model of the standard deviation within each replicate group resulted in a standard deviation of 0.085 (Supplementary File Figure 1) of the cycle threshold. The 95% confidence interval of the difference of the cycle threshold was therefore determined as 0.24 (2*[√2 * 0.085]) , and this value was used for the removal of outlier technical replicate CT values. CT values within each technical replicate group were therefore removed from the dataset when falling outside of this confidence interval, removing those samples with the largest standard deviation (Supplementary File Figure 2).

Following the removal of outlier deltaCT values, the final datasets for differential gene expression analysis included 578 of the original 607 deltaCT values for snail line comparisons (Table 3, Supplementary Table 2, Supplementary File Figure 3) and 256 of 259 deltaCT values for diet comparison (Table 4, Supplementary Table 3, Supplementary File Figure 4).

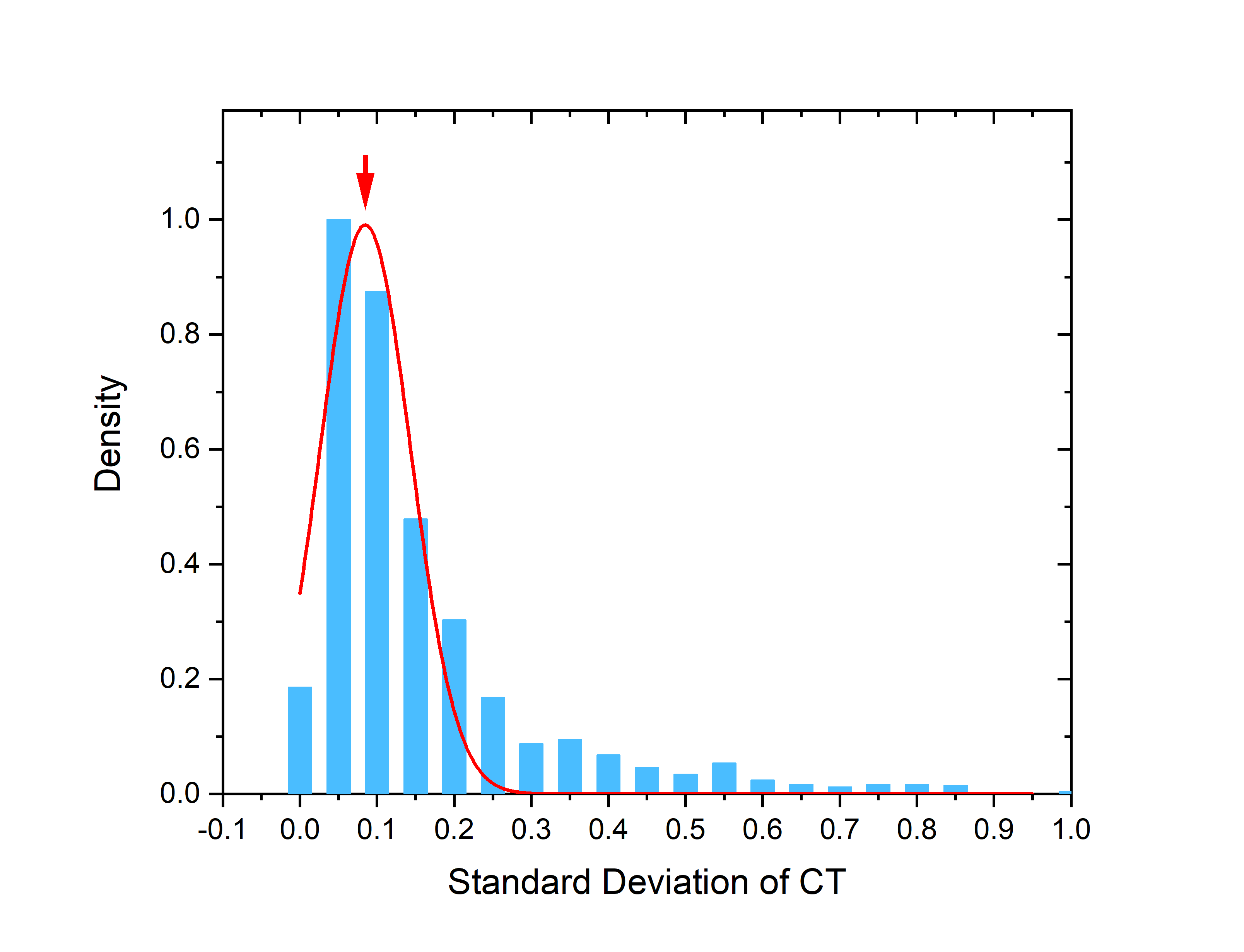

Supplementary Figure 1. Histogram of the standard deviation of cycle threshold calculated within each replicate group (i.e. between the three replicate datapoints reported per sample, per gene) showing the Gaussian fitted model with a prominent peak at 0.085.

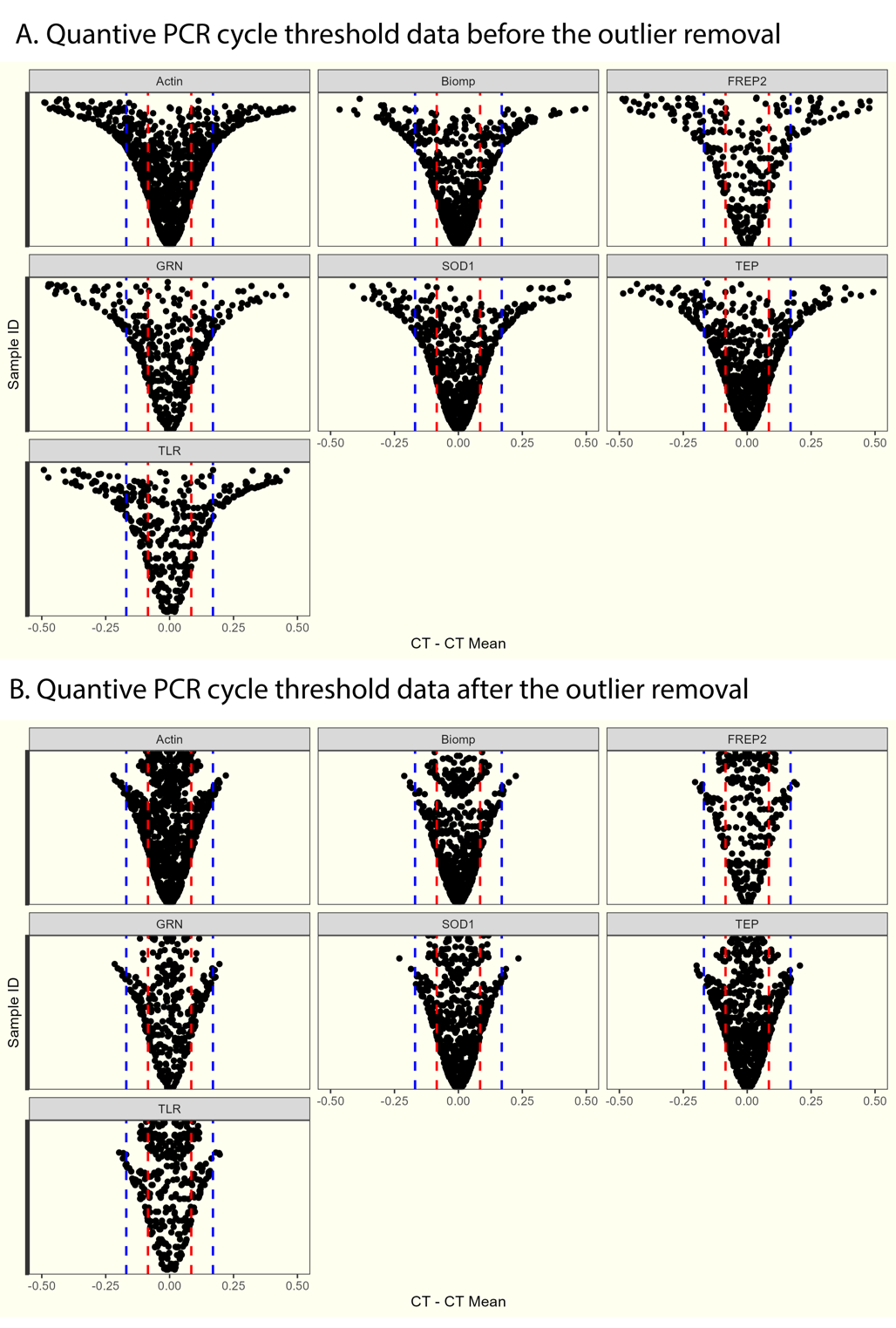

Supplementary Figure 2. The variation of the qPCR cycle threshold (CT) data in the current study before (A) and after (B) the removal of outliers within each replicate group (within each sample and each gene target). The x-axis represents the replicates CT value minus the average CT of the corresponding replicate group, and the y-axis represents each sample sorted by the standard deviation of each replicate group. Red dashed lines indicate the plus and minus one standard deviation of the qPCR data (0.085), and the blue lines indicate the plus and minus two standard deviation (0.17). It is shown that the samples with the largest standard deviations were removed after the process, and those that lie outside of the 95% confidence interval of +/- 0.24 CT.

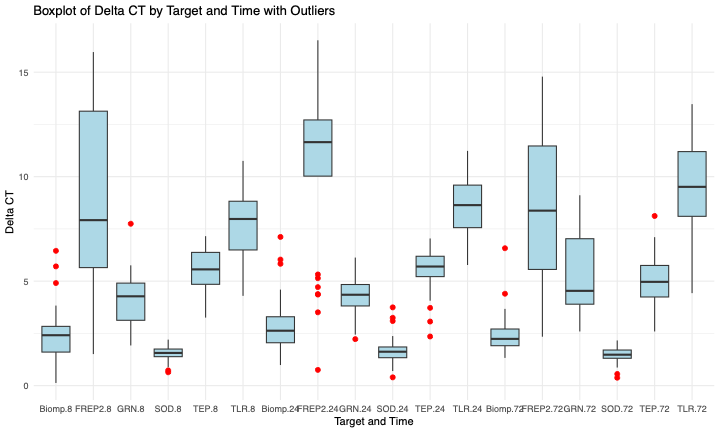

**Supplementary Figure 3.** Boxplot of the delta CT values for the five target genes (FREP2 not included in this study) and three time points when comparing gene expression in *Biomphalaria sudanica* line 163 and KEMRIwu exposed to *S. mansoni* or sham exposed, with outlier delta CT values (below 25^th^ percentile or above 75^th^ Percentile) that were removed prior to statistical analysis.

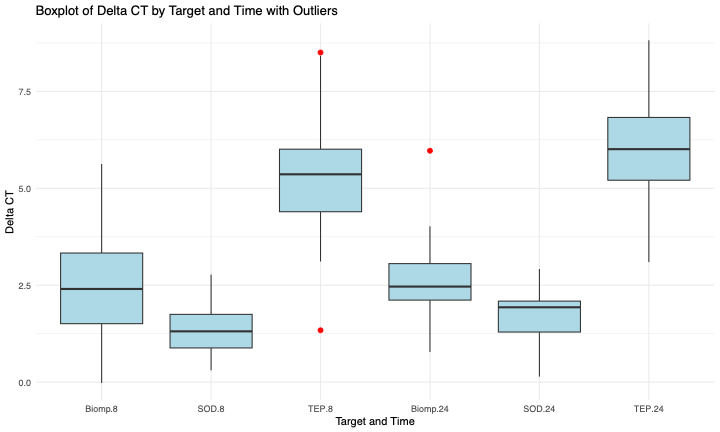

**Supplementary File Figure 4.** Boxplot of the deltaCT values for three target gene and two time points when comparing gene expression in *Biomphalaria sudanica* line KEMRIwu fed either a strict diet of green leaf lettuce or a commercial food pellet exposed to *S. mansoni* or sham exposed, with outlier deltaCT values (below 25^th^ percentile or above 75^th^ Percentile) that were removed prior to statistical analysis.
